# RNetDys: identification of disease-related impaired regulatory interactions due to SNPs

**DOI:** 10.1101/2022.10.08.511312

**Authors:** Céline Barlier, Mariana Messias Ribeiro, Sascha Jung, Antonio del Sol

## Abstract

The dysregulation of regulatory mechanisms due to Single Nucleotide Polymorphisms (SNPs) can lead to diseases and does not affect all cell (sub)types equally. Current approaches to study the impact of SNPs in diseases lack mechanistic insights. Indeed, they do not account for the regulatory landscape to decipher cell (sub)type specific regulatory interactions impaired due to disease-related SNPs. Therefore, characterizing the impact of disease-related SNPs in cell (sub)type specific regulatory mechanisms would provide novel therapeutical targets, such as promoter and enhancer regions, for the development of gene-based therapies directed at preventing or treating diseases. We present RNetDys, a pipeline to decipher cell (sub)type specific regulatory interactions impaired by disease-related SNPs based on multi-OMICS data. RNetDys leverages the information obtained from the generated cell (sub)type specific GRNs to provide detailed information on impaired regulatory elements and their regulated genes due to the presence of SNPs. We applied RNetDys in five disease cases to study the cell (sub)type differential impairment due to SNPs and leveraged the GRN information to guide the characterization of dysregulated mechanisms. We were able to validate the relevance of the identified impaired regulatory interactions by verifying their connection to disease-related genes. In addition, we showed that RNetDys identifies more precisely dysregulated interactions linked to disease-related genes than expression Quantitative Trait Loci (eQTL) and provides additional mechanistic insights. RNetDys is a pipeline available at https://github.com/BarlierC/RNetDys.git

## Introduction

Gene regulation is largely controlled by the binding of transcription factors (TFs) to regulatory elements, such as promoters and enhancers, to control cell (sub)type specific functions. Notably, it has been shown that most of these functions are strongly regulated by enhancer activity (Latchman, 2011; Andersson et al., 2014). Therefore, the impairment of the regulatory interactions between TFs and enhancers of regulated genes can lead to dysregulations that trigger pathological gene expression changes that contribute to disease development (Lee and Young, 2013). In that regard, Single Nucleotide Polymorphisms (SNPs) have been shown to be associated with regulatory dysregulations driving complex diseases, such as diabetes and Alzheimer’s disease (AD) (Hiramoto et al., 2015; Akhlaghipour et al., 2022). Standard approaches such as Genome-Wide Association Studies (GWAS) and expression Quantitative Trait Loci (eQTLs) have been used to study the association between SNPs and genes (Visscher et al., 2017; Bryois et al., 2022; Gazal et al., 2022). In particular, GWAS successfully deciphered thousands of disease-related SNPs (Claringbould and Zaugg, 2021). Moreover, GWAS showed that the majority of these SNPs were found in non-coding regions, particularly in enhancer regions, and thus were most likely involved in gene regulation (Nica and Dermitzakis, 2013). Moreover, eQTLs have been useful to provide further insights in understanding the influence of SNPs in diseases by associating them to their target genes, based on the statistical association of gene expression variation to these genetic polymorphisms (Jeng et al., 2020). However, these approaches only provide information on SNP-gene relationships. Leveraging multi-OMICS data to construct and exploit the regulatory landscape in order to gather additional mechanistic insights would significantly contribute to a better understanding of the impact of disease-related SNPs on gene regulation and disease development. Notably, GRNs have been widely used to gain insights into diseases (Emmert-Streib et al., 2014; Ament et al., 2018; Bakker et al., 2021) but the characterization of underlying regulatory mechanisms dysregulated due to SNPs and the cell (sub)types specifically impaired remains elusive. The resolution of cell (sub)type specific regulatory mechanisms impaired due to SNPs in disease would provide additional mechanistic insights and pave the way towards the development of gene-based therapies for disease prevention and treatment (Uddin et al., 2020).

We present RNetDys, a multi-OMICS pipeline that identifies impaired regulatory mechanisms due to the presence of disease-related SNPs at the cell (sub)type level. In particular, RNetDys combines scRNA-seq, scATAC-seq, ChIP-seq and prior-knowledge to build comprehensive cell (sub)types or state specific GRNs that are leveraged to capture impaired interactions due to disease-related SNPs. Compared to existing strategies to study SNPs (Farh et al., 2014, Yu et al., 2022; Nathan et al., 2022), this pipeline provides a comprehensive view of the impaired regulatory landscape, including interactions mediated by TFs and enhancers of regulated genes and activation or repression mechanisms to provide additional mechanistic insights. In particular, RNetDys provides the binding affinity score of impaired TFs, the type of mechanism dysregulated, and a list of ranked TFs based on their importance in the impaired network topology, the strength of the binding impairment and the frequency of SNPs occurring in the global population.

We applied RNetDys in five disease case studies and showed that it was able to accurately capture impaired regulatory interactions and provide additional mechanistic insights by leveraging the information obtained from the GRN inference.

## Material and methods

### General workflow of RNetDys

We implemented a systematic pipeline integrating different type of OMICS data to decipher impaired regulatory mechanisms due to SNPs in disease by leveraging the GRN information. The pipeline was divided in two main parts composed of the cell (sub)type specific GRN inference and the capture of impaired regulatory interactions due to disease-related SNPs to gain regulatory mechanistic insights for the disease.

#### Cell (sub)type specific regulatory interactions inference

The cell (sub)type specific regulatory network inference was based on a multi-OMICS approach that used single cell transcriptomics and single cell chromatin accessibility, not necessarily matched, as well as prior-knowledge, including ChIP-seq data and reported enhancers interactions. First, using the scRNA-seq we selected genes that were conserved at least in 50% of the cells for further analyses. Then, we ensured the accessibility of the corresponding promoter regions using scATAC-seq data and predicted TF-promoter interactions by intersecting the ChIP-seq TF-binding evidence with the open promoter regions using BEDTools (Quinlan and Hall, 2010). Then, we performed a peak correlation using the scATAC-seq data and carried out a statistical test, as well as a BH multiple correction, to select the significant interactions such as p-adjusted value < 0.05. The identified enhancer-promoter interactions were then intersected with GeneHancer (Fishilevich et al., 2017), used as a backbone and interactions involving active promoters were kept. Then, TF-enhancers interactions were inferred by intersecting the ChIP-seq and scATAC-seq data. Finally, the regulatory interactions were signed to distinguish activations from repressions by computing the Pearson correlation between TFs and genes using the scRNA-seq dataset (Fig. S1). Correlation scores for enhancer-promoter interactions were computed such as:

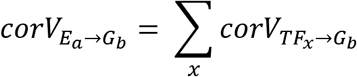

With corV corresponding to the correlation value, E denoting the enhancer and G corresponding to the gene. And, correlation scores for TF-enhancer were computed such as:

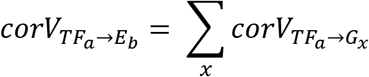

With corV denoting the correlation value, E corresponding to the enhancer and G to the gene. Finally, positive correlation scores were considered to be activations whereas negative ones were considered to be repressions. Further details are provided in Supplementary Information.

#### Identify candidate impaired regulatory interactions

Using the cell (sub)type specific GRN inferred in healthy condition, we then contextualized the GRN towards the disease condition. The contextualization required a list of SNPs for the disease studied and the cell (sub)type GRN of interest. The SNPs were mapped to the GRN by using their coordinates and interactions for which a SNP was falling into a TF binding region of an enhancer or promoter were considered as candidates to be impaired in the dis-ease. We then performed a TF binding analysis using PERFECTOS-APE (E. Vorontsov et al., 2015) to refine the candidate interactions by selecting the ones having at least one binding site significantly impaired by the SNP (Supplementary Information). Finally, we ranked TFs by their involvement in the regulatory impairments based on the network topology and the MAF score of SNPs such as:

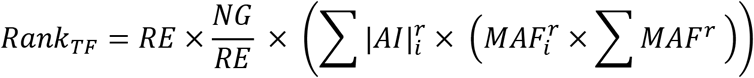

Where RE denote the number of regulatory elements regulated by the TF, NG corresponds to the number of downstream genes across RE, AI denotes the binding affinity impairment log2FC and i corresponds to the SNPs and r the regulatory element.

### Prior-knowledge collection and processing

RNetDys relied on prior-knowledge data that were collected and processed to be integrated in the pipeline. The ChIP-seq bed files were downloaded from ChIP Atlas (Oki et al., 2018) for human hg19 and hg38 assemblies. Bed files were annotated using HOMER (Heinz et al., 2010) with the latest GTF file for each assembly. Enhancer regions and their connected genes were obtained from the GeneHancer database (Fishilevich et al., 2017). Of note, GeneHancer database provided information for hg38 coordinates and hence, we used LiftOver (https://genome.ucsc.edu/cgi-bin/hgLiftOver) to convert these coordinates for hg19 to provide more flexibility to our pipeline.

### Data collection and analysis

First, to perform the benchmarking analysis, we collected 20 publicly available scRNA-seq and 11 scATAC-seq datasets from six human cell lines including BJ, GM12878, H1-ESC, A549, Jurkat and K562 (Table S1). Then, we collected scRNA-seq and scATAC-seq healthy data from pancreas and brain tissues to extract cell (sub)types using Seurat (Hao et al., 2021) and Signac (Stuart et al., 2020), and then generated the GRNs (Supplementary information). Finally, we collected SNPs from ClinVar (Landrum et al., 2018) for five diseases including Alzheimer’s disease (AD), Parkinson’s disease (PD), Epilepsy (EPI), Diabetes type I (T1D) and type II (T2D) to perform the network contextualization towards the disease condition. Notably, SNPs were defined as being single nucleotide variants found at least in 1% of the global population such as MAF >= 0.01 (Supplementary Information). In addition, we performed an outdegree analysis for three main TFs involved in the regulatory impairments. The outdegree ratios were computed by scaling each TF outdegree by the maximum outdegree in each cell (sub)type.

### Validation and comparison to state-of-the-art

We first assessed the performances of RNetDys in identifying cell (sub)type specific regulatory interactions and compared them to state-of-the-art GRN inference methods (Aibar et al., 2017; Chan et al., 2017; Kim, 2015; Huynh-Thu et al., 2010) (Supplementary Information). First, we benchmarked the performances of each method to infer cell (sub)type specific TF-promoter interactions. The gold standards (GS) were compiled using cell line specific ChIP-seq from Cistrome (Mei et al., 2017) by selecting only the highest quality data. Then, we assessed the performances of RNetDys for capturing cell (sub)type specific enhancer-promoter regulatory interactions compared to Cicero, a widely used method to identify cis-interactions based on scATAC-seq data (Pliner et al., 2018). The GS networks were built using promoter capture Hi-C data from 3DIV (Yang et al., 2018) for three of the human cell lines. For both benchmarking analyses, we computed the precision (PPV) and F1-score (F1) to assess the performances such as:

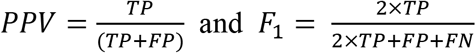

With TP = True Positive (predicted and found in the GS), FP = False Positives (predicted but not in the GS) and FN = False Negatives: (not predicted but in the GS).

We then compared the ability of RNetDys to precisely capture gene-disease relationships in cell (sub)types, compared to eQTL (Bryois et al., 2022). First, we downloaded Online Mendelian Inheritance in Man (OMIM) Morbid Map (Amberger et al., 2019), filtered for gene-disease interactions reported in the five diseases in study (AD, EPI, PD, T1D, and T2D), and removed the interactions reported as provisional. Then, we matched these gene interactions to the SNP-associated genes identified by eQTL and RNetDys. The ratio of matched genes in eQTL and RNetDys was calculated by dividing the number of matched genes by the total of genes identified in each of the methods across all cell (sub)types.

## Results

### RNetDys, a multi-OMICS pipeline to decipher impaired regulatory mechanisms

We implemented RNetDys, a systematic pipeline based on multi-OMICS data to decipher impaired regulatory interactions due to SNPs in diseases by leveraging the information of cell (sub)type specific GRNs (Fig.1).

**Fig. 1.**
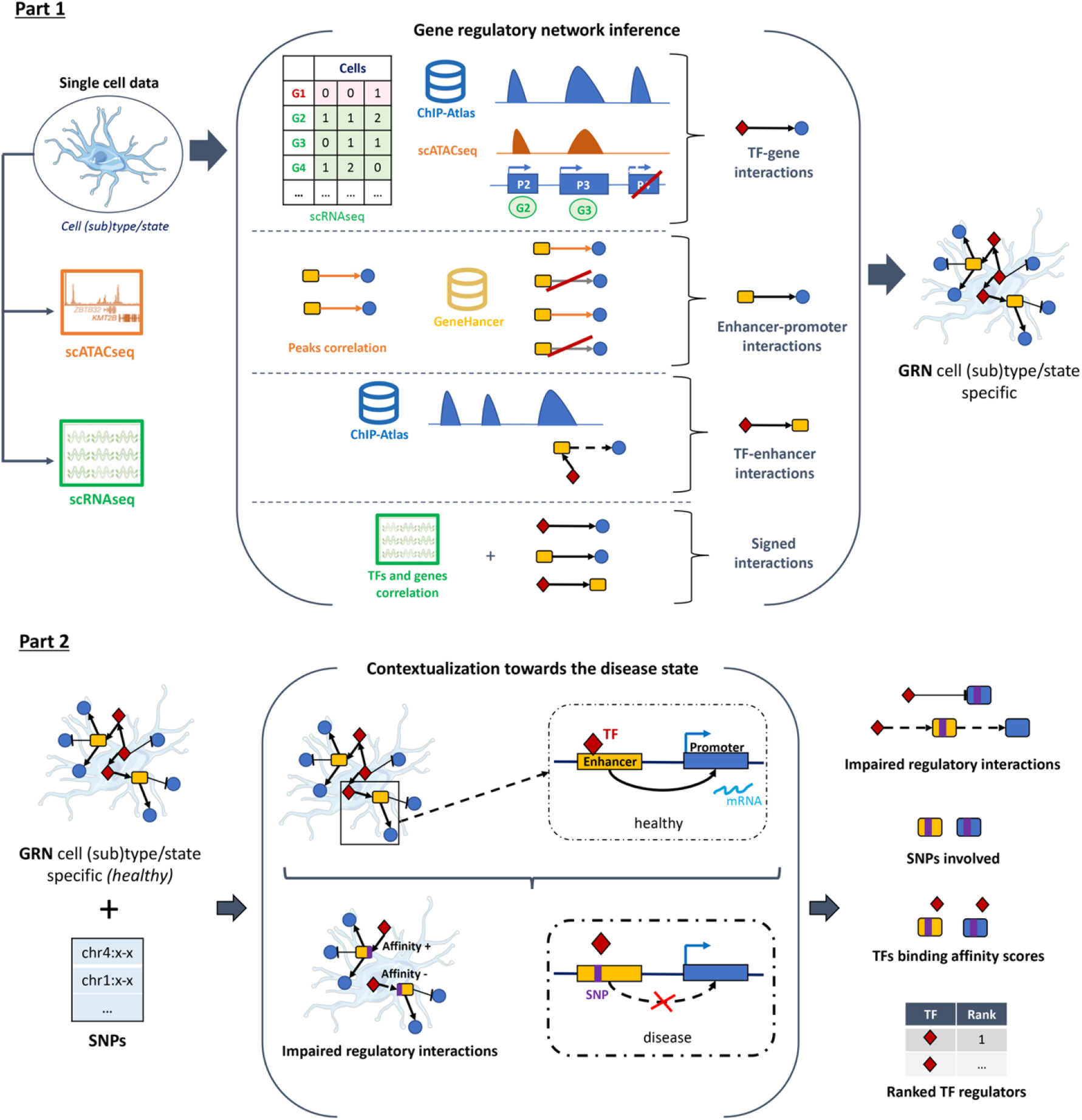
General workflow of RNetDys to decipher regulatory dysregulation in diseases. RNetDys is composed of two main parts including (1) the cell (sub)type specific GRN inference using scRNA-seq, scATAC-seq and prior-knowledge, and (2) the identification of candidates impaired regulatory interactions using the GRN and a list of SNPs, followed by the TF-binding affinity analysis. The part one provides the cell (sub)type or state specific GRN describing the regulatory interactions mediated by TFs and enhancers of regulated genes. The part two provides the list of candidate impaired regulatory interactions in the cell (sub)types, the SNPs that were mapped to these interactions and the TFs for which the binding ability might be impaired and regulatory TFs ranked based on their importance in the impairments.

RNetDys is an integrative approach relying on single cell transcriptomics and single cell chromatin accessibility from a specific cell (sub)type or state, as well as prior-knowledge information including extensive ChIP-seq data (Oki et al., 2018) and reported enhancer-promoter relationships (Fishilevich et al., 2017). The pipeline is composed of two main parts: (i) the cell (sub)type specific GRN inference and (ii) the identification of impaired regulatory mechanisms due to SNPs in diseases (Fig. 1, Fig. S1). The first part consists of the GRN inference for a healthy cell (sub)type or state based on scRNA-seq and scATAC-seq data as an input. Notably, the two single cell datasets do not need to be matched but they need to contain the same cell (sub)type. The second part takes as an input a cell (sub)type or state specific GRN and a list of SNPs of particular interest for the disease studied (Visscher et al., 2017; Landrum et al., 2018). In particular, the SNPs provided could have been described as related to the disease of interest in prior-knowledge databases (Landrum et al., 2018) or identified by genotyping analyses (Nielsen et al., 2011). As a result, RNetDys provides the impaired regulatory mechanisms, the corresponding SNPs, the affinity scores of TF having their binding site impaired, and a list of ranked TF regulators based on their involvement in the observed impairments (Fig. 1).

### RNetDys is more accurate to infer cell (sub)type specific GRNs

RNetDys mainly relies on the cell (sub)type specific regulatory landscape to identify impaired regulatory interactions due to disease-related SNPs. Therefore, we assessed the performance of RNetDys in predicting cell (sub)type specific GRNs (Fig. 2). In this regard, we performed the benchmarking of both TF-gene and enhancer-promoter interactions, compared to current methods. We showed that our approach overcame the state-of-the-art GRN inference methods for predicting cell (sub)type specific TF-gene interactions with an average precision of 0.20 and average accuracy of 0.28 (Fig. 2A, B).

**Fig. 2.**
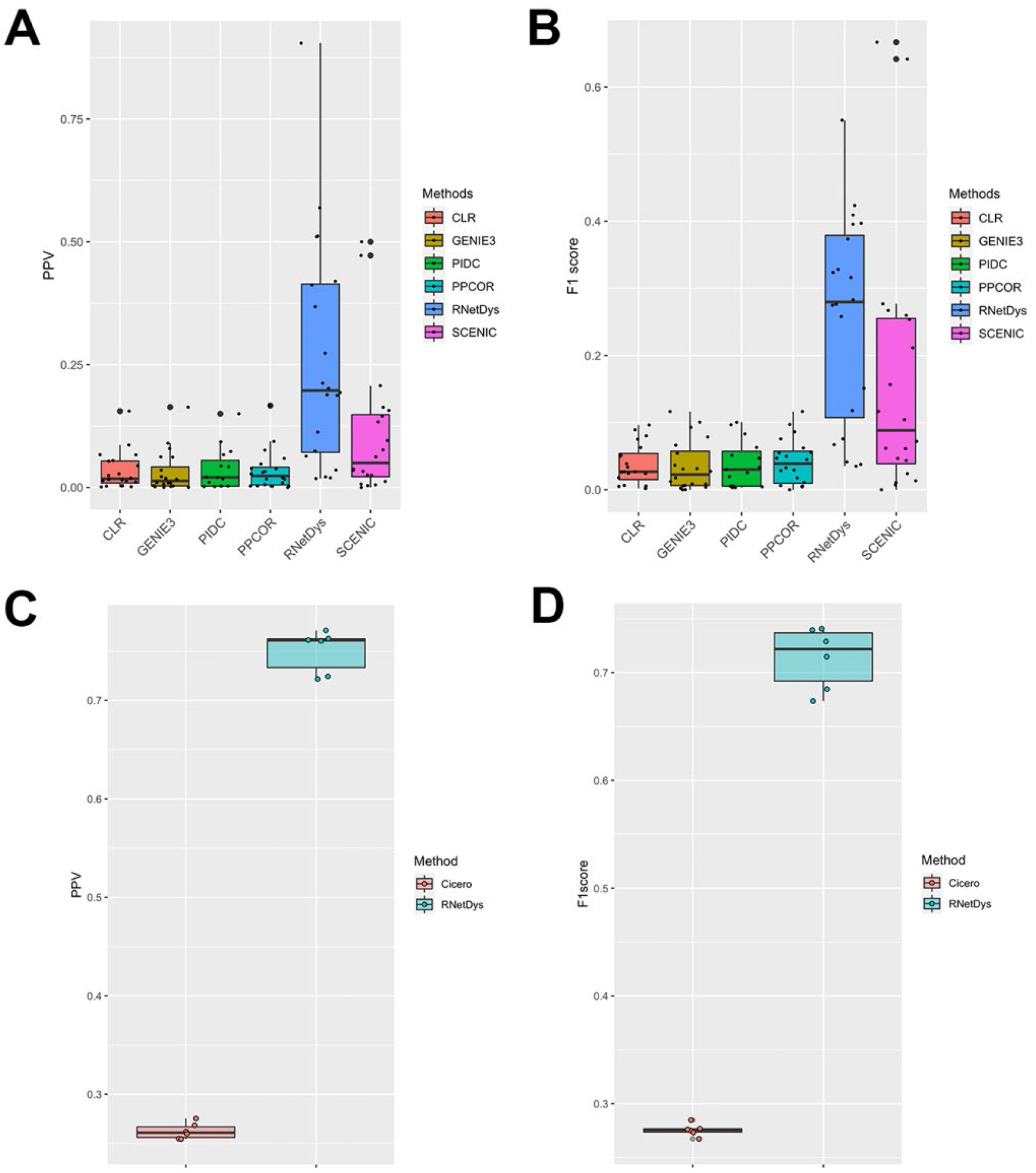
Performances of RNetDys and comparison to other methods. (A, B) TF-promoter regulatory interactions performances assessed using (A) the PPV and (B) the F1-score metrics. Performances were assessed for RNetDys, state-of-the-art methods and metrics on 20 datasets from six human cell lines. (C, D) Enhancer-promoters regulatory interactions performance assessment using (C) the PPV and (D) the F1-score metrics. Performances were assessed for RNetDys and Cicero on 6 scATAC-seq datasets from three human cell lines.

This assessment highlighted the strength of combining different regulatory layers with prior-knowledge to provide predictions with a higher confidence. Moreover, we showed that RNetDys outperformed Cicero in capturing cell (sub)type specific enhancer-promoter interactions with a median precision of 0.76 and median accuracy of 0.72, supporting the confidence provided by the prior-knowledge leveraged by our approach (Fig. 2C, D). This analysis demonstrated the accuracy of the cell (sub)type specific GRN information leveraged by our pipeline to capture impaired transcriptional regulatory mechanisms due to SNPs in diseases.

### RNetDys provides additional insights into the mechanistic dysregulation enhanced by SNPs

#### Validation of the SNPs impairment and comparison to state-of-the-art approaches

We applied RNetDys to five diseases, including AD, PD, EPI, T1D and T2D, by collecting disease-related SNPs from ClinVar (Landrum et al., 2018) and cell (sub)type specific GRNs generated from human pancreas and brain tissues. First, we supported the relevance of predicted SNP-gene interactions identified by RNetDys using available GWAS data from ClinVar database and recently published cell-type specific eQTL information (Bryois et al., 2022). Across the five diseases, we were able to find support for 90% of the SNP-target gene relationships identified by our pipeline (Table S4). Furthermore, by using cell type specific eQTL data, we were able to validate the occurrence of certain SNPs and their impact on the predicted target genes in specific cell types. For instance, our results show that the PD-associated SNPs rs11538371, rs2072814 and rs8137714 are found to be linked to TIMP3 in astrocytes (Table S4). In fact, TIMP3 is an inhibitor of metalloproteinases, enzymes secreted by astrocytes (Yin et al., 2006), that are implicated in several PD-associated processes such as dopaminergic neuron degeneration, neuroinflammation, and proteolysis of α-synuclein (Sung et al., 2005; Choi et al., 2011; Annese et al., 2015). Second, we evaluated the precision of RNetDys in capturing gene-disease relationships at the cell(sub)type level using the OMIM database (Amberger et al., 2019) (Fig. S2). When compared to the eQTL data, we observed that the genes captured by our approach as being impaired due to the presence of SNPs are more often linked to disease than the genes captured by eQTL. Although eQTL captures a larger number of SNP-gene interactions, few of them are actually described to be involved in the disease, thus explaining the low ratio. On the other hand, RNetDys identifies more genes linked to each SNP that have been described as related to the disease, demonstrating the higher precision of RNetDys compared to eQTL.

#### Cell (sub)type differential dysregulation in diseases

Then, we studied the differential impairment across cell (sub)types in the five diseases as it has been reported that some cell (sub)types were more involved in disease mechanisms (Muratore et al., 2017; Kamath et al., 2022). We observed that cell (sub)types shared few impaired interactions in the studied diseases, especially in EPI and PD (Fig. 3). Interestingly, in EPI, astrocytes, OPCs and inhibitory neurons seem to be the most impaired cell types. This is consistent with literature evidence that shows that modifications in GABA receptors, which are expressed in inhibitory neurons, are closely linked to epilepsy (Tanaka et al., 2012). Furthermore, impairment of antiquin expression, encoded by the gene ALDH7A1, in astrocytes has been described to be linked with dysregulation of neurotransmitter shuttling and recycling, one of the major causes of neurological deficits (David et al., 2009; Jansen et al., 2014). Finally, studies showed that myelinated neuronal axons are damaged in epileptic patients and the ability of OPCs to proliferate is reduced in samples obtained from patients with dysplasia (Luo et al., 2015; Donkels et al., 2020).

**Figure 3.**
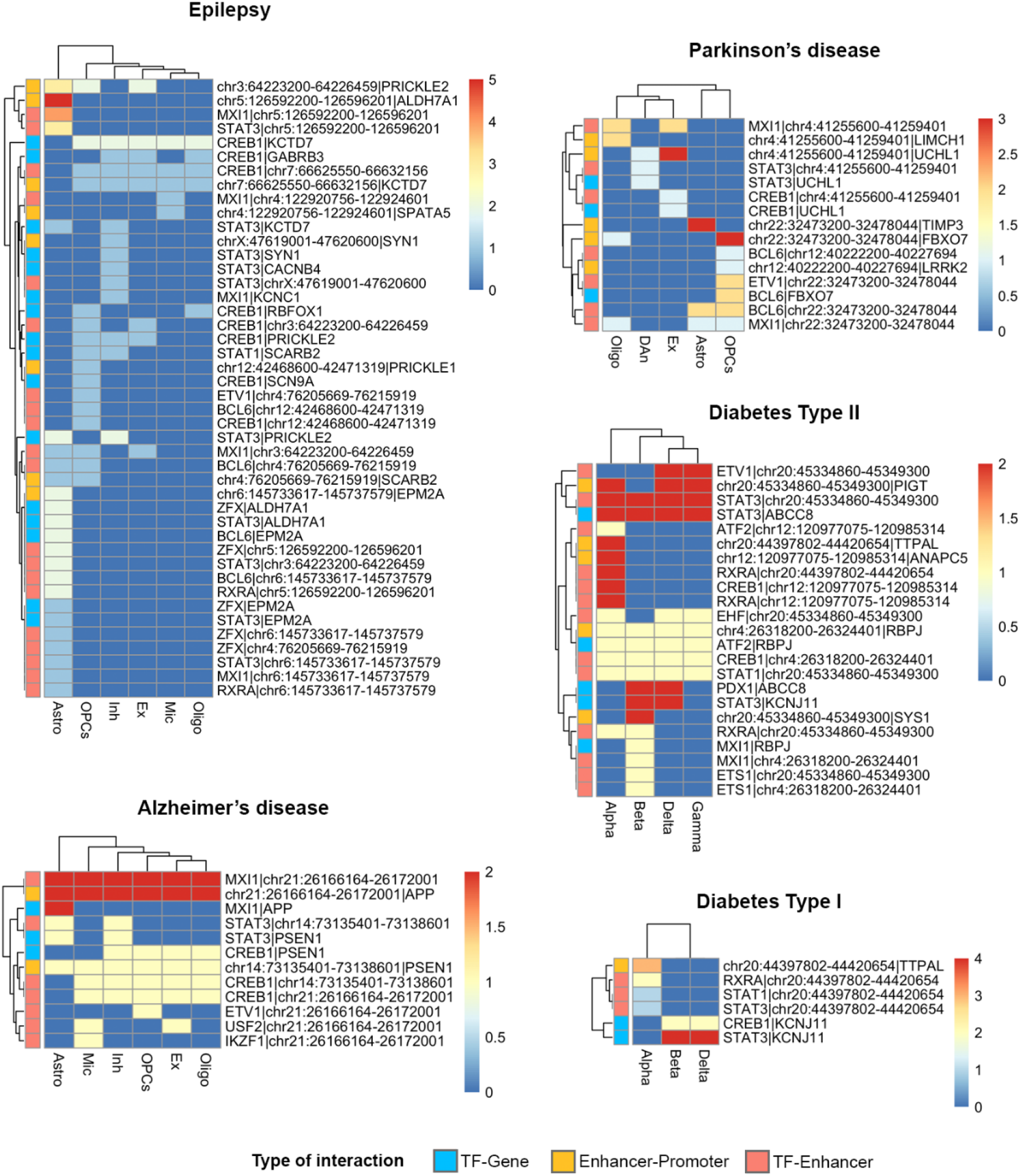
Cell (sub)type differential regulatory impairment in diseases. Heatmaps showing the distribution of impaired interactions due to disease-related-SNPs across cell (sub)types for Alzheimer’s disease (AD), Parkinson’s disease (PD), Epilepsy (EPI), Diabetes type I (T1D) and type II (T2D). The colors of the heatmap represent the number of SNPs impacting the regulatory interactions. Astro: astrocytes, Ex: excitatory neurons, Inh: inhibitory neurons, Mic: microglia, Oligo: oligodendrocytes, OPCs: oligodendrocyte progenitors, DAn: dopaminergic neurons.

#### Insights into the cell (sub)type specific regulatory impairments

We finally aimed at exploiting the GRN information provided by RNetDys to further analyse the regulatory impairments of cell (sub)types (Fig. 4, Fig. S3-S6). We observed that in AD (Fig. 4), the same enhancers were involved in all cell (sub)types specific networks with an impact on the expression of APP and presenilin 1 (PSEN1). Alterations in the expression of these genes are primarily linked to the development of AD (Dewachter et al., 2002; Matsui et al., 2007). Furthermore, recent studies have shown that not only neurons, but also astrocytes and microglia to be involved in the accumulation of β-amyloid plaques (Palop and Mucke, 2010; Frost and Li, 2017). However, the impairment of the TFs and enhancers regulating these two genes seems to be different across cell (sub)types (Fig. 4). Indeed, most of the SNPs in astrocytes and microglia would induce a repression of APP whereas this gene seems to be activated in other cell (sub)types (Fig. 4). It has been described that these two cell types provide protective effects, with microglia facilitating the clearance of β-amyloid overproduced by neurons in AD (Fakhoury, 2018).

**Figure 4.**
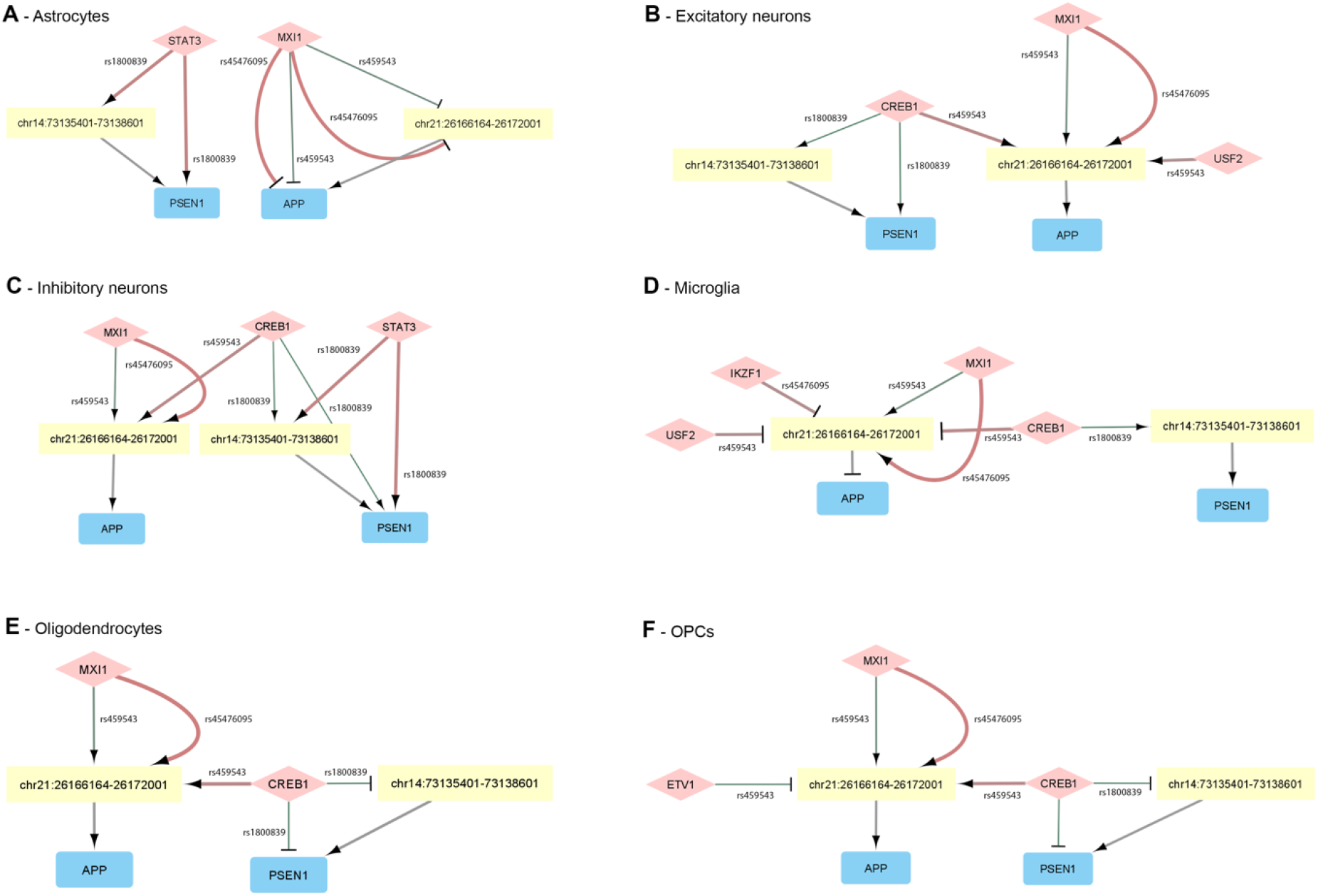
Cell (sub)type specific regulatory impairment in AD. Network visualization of impaired regulatory interactions for (A) astrocytes, (B) excitatory neurons, (C) inhibitory neurons, (D) microglia, (E) oligodendrocytes and (F) OPCs. TFs are represented as diamonds in light red, enhancers as yellow rectangles and genes in blue rectangles. Arrows represent activations and T edges represent repressions. The weight of edges from TFs correspond to the strength of the impairment, with the thinnest translating a weaker binding affinity and a large edge being a strong increase in binding affinity. The color of the edges from TFs represents the log2FC with green being a decreased affinity and red an increased one.

To provide better insights on the main regulatory TFs behind disease dysregulation, we ranked the impaired TFs based on network topology and impact of each involved SNP (see Methods). Notably, we could observe that certain TFs, such as CREB1, MXI1, and STAT3, are ranked as top regulators across different brain diseases and diabetes (Table 1).

**Table 1.** TF regulators involved in impaired regulatory mechanisms.

| **DISEASE** | **CELL (SUB)TYPE** | **RANKED TFS\*** |
| --- | --- | --- |
| AD | Astrocyte | MXI1, STAT3 |
|  | Excitatory neuron | CREB1, USF2, MXI1 |
|  | Inhibitory neuron | CREB1, MXI1, STAT3 |
|  | Microglia | CREB1, USF2, MXI1, IKZF1 |
|  | Oligodendrocyte | CREB1, MXI1 |
|  | OPCs | CREB1, MXI1, ETV1 |
| EPI | Astrocyte | MXI1, STAT3, BCL6, ZFX, RXRA |
|  | Excitatory neuron | CREB1, MXI1 |
|  | Inhibitory neuron | CREB1, STAT3, STAT1, MXI1 |
|  | Microglia | CREB1, MXI1 |
|  | Oligodendrocyte | CREB1 |
|  | OPCs | CREB1, BCL6, MXI1, STAT1, ETV1 |
| PD | Astrocyte | MXI1, BCL6 |
|  | Dopaminergic neuron | STAT3 |
|  | Excitatory neuron | MXI1, CREB1 |
|  | Oligodendrocyte | MXI1 |
|  | OPCs | BCL6, MXI1, ETV1 |
| T1D | Alpha cell | STAT3, STAT1, RXRA |
|  | Beta cell | STAT3, CREB1 |
|  | Delta cell | STAT3, CREB1 |
| T2D | Alpha cell | STAT3, RXRA, STAT1, CREB1, ATF2, EHF |
|  | Beta cell | CREB1, STAT1, STAT3, PDX1, ETS1, ATF2, RXRA, MXI1 |
|  | Delta cell | CREB1, STAT1, STAT3, PDX1, ETV1, EHF, ATF2 |
|  | Gamma cell | STAT3, CREB1, STAT1, ETV1, EHF, ATF2 |

\* TFs are ranked by their order of importance in the detected impaired regulatory mechanisms.

To investigate this, we evaluated the outdegree distribution of these TFs and we observed different outdegrees values across cell (sub)types, which demonstrates that our ranking has no bias towards highly connected TFs (Fig. S7). CREB1, MXI1, and STAT3 participate in common cell mechanisms involved disease development, such as cell death and inflammation. However, each of these TFs has been described to play a different function in these mechanisms in different diseases.

For instance, MXI1 has been shown to be involved in the aging of the neurovascular unit, which contributes for the progression of AD (Zhao et al., 2022). On the other hand, the same TFs seems to be part of the unique transcriptomic signature of T2D, which we can also observe in our results as this TF does not show as a key regulator in T1D (Table 1) (Cubillos-Angulo et al., 2020). Finally, MXI1 was found to be one of the main regulators involved in impaired regulatory interactions for PD, apart from dopaminergic neurons (Fig. S3). MXI1 has been described to be involved in the mitochondrial homeostasis, dysregulated in PD and known to be involved with neurodegeneration (Lestón Pinilla et al., 2021; Malpartida et al., 2021).

CREB1 has been extensively shown to regulate gluconeogenesis through the coactivator PGC-1, playing a vital role in the regulation of efficient glucose sensing and insulin exocytosis and in the development of diabetes (Herzig et al., 2001). Our results show this TF to be the main regulator involved in AD and EPI in all cell (sub)types, apart from astrocytes (Table 1, Fig. 4, Fig. S4). CREB1 is a TF responsible for regulating the major pathways that mediate neurotrophin-associated gene expression, a group of proteins that promotes survival and neuronal development (Shaywitz and Greenberg, 1999). Indeed, increased CREB activity promotes hyperexcitability, one of the main triggers of seizures, while reduced levels seem to prevent epilepsy (Zhu et al., 2012; Wang et al., 2020) (Fig. S4). PSEN1 has been shown to be a downstream target of CREB1 (Cui et al., 2022), which further supports the results obtained by our pipeline as CREB1 was predicted to regulate PSEN1. In AD, PSEN1 upregulation leads to myelin dysfunction in OPCs in cases of familial AD (Desai et al., 2011). Notably, our pipeline predicts a decrease in CREB1 binding affinity to the promoter and enhancer regions of PSEN1 in the presence of rs1800839, potentially elucidating one of the possible mechanisms behind PSEN1 upregulation previously observed in AD (Fig. 4).

Finally, STAT3 was overall found to be the main regulator involved in impaired interactions of T1D and T2D (Table 1, Fig.s S5 and S6). In the pancreas, STAT3 has been shown to regulate insulin secretion and islet development (Saarimäki-Vire et al., 2017). In addition, in T2D, exacerbated STAT3 signalling has been shown to lead to insulin resistance in skeletal muscle of diabetic patients (Mashili et al., 2013), supporting its importance as a regulator of the dysregulations involved in the disease. In neurodegenerative diseases, STAT3 activation has been shown to promote astrogliosis, which is reflected in our results by an increase of the binding affinity of this TF to distinct regulatory regions (Fig. 4A and Fig. S4A) (Toral-Rios et al., 2020).

## Discussion

The study of cell (sub)type or state specific regulatory interactions impaired due to disease-related SNPs is required to pave the way towards the development of gene-based therapies to prevent or treat diseases (Rao et al., 2021). In addition, the comprehensive view of the regulatory landscape, including interactions mediated by TFs and enhancers of regulated genes, is critical to study dysregulated mechanisms in diseases (Emmert-Streib et al., 2014; Chiou et al., 2021). In that regard, existing strategies to study the impact of SNPs do not exploit the GRN information to provide additional mechanistic insights into the disease-related dysregulations (Rao et al., 2021; Bryois et al., 2022). In addition, recent studies have shown that specialized group of cells, including cell types, subtypes and phenotypes, are not equally involved in diseases (Nathan et al., 2022; Kamath et al., 2022). However, current approaches have been mainly focused on cell types, lacking ability to identify dysregulated mechanisms at deeper levels of resolution. RNetDys is a systematic multi-OMICS pipeline to decipher cell (sub)type or state specific regulatory interactions impaired due to SNPs in diseases. This pipeline exploits the high-resolution of single cell to infer a comprehensive regulatory landscape, leveraged to identify impairment due to SNPs. We applied RNetDys to five disease cases and showed that cell (sub)types specific regulatory mechanisms were not equally impaired, suggesting their differential involvement in the studied diseases. Moreover, we validated the relevance of some impaired regulatory mechanisms using GWAS and eQTL data (Landrum et al., 2018; Bryois et al., 2022). In that regard, we provided additional mechanistic insights into the regulatory mechanisms dysregulated and identified the main TF regulators involved. Notably, the presented analysis was performed using SNPs retrieved from ClinVar, but RNetDys could be of great use to provide valuable regulatory mechanistic insights by using SNPs derived from genotyping studies. In the present study, we were able to predict known and unreported cell (sub)type specific SNP-gene interactions, hence showing how our pipeline could facilitate the discovery of regulatory impairments. To conclude, we foresee RNetDys to be a valuable tool to comprehensively identify cell (sub)type specific regulatory mechanisms impaired due to SNPs and aid the development of strategies for therapeutic intervention in diseases.

## Data and Material availability

RNetDys is a pipeline publicly available at https://github.com/BarlierC/RNetDys.git.

The repository of generated regulatory networks, results and scripts used in this study are available at https://gitlab.com/C.Barlier/RNetDys_analyses.

## Acknowledgements

The authors thank Dr. Patrick May for the valuable feedback and insights provided for this project. The benchmarking of the state-of-the-art GRNs method and data processing was performed using the HPC facilities of the University of Luxembourg (https://hpc.uni.lu).

## Author contributions

C.B. implemented RNetDys, collected and processed the data, performed the benchmarking of the GRNs, generated the cell (sub)type specific GRNs, collected the disease-related SNPs, performed the data analysis and wrote the manuscript, M.M.R. collected and processed the data, extracted the healthy cell (sub)types datasets, performed the validation of impaired interactions, the data analysis and wrote the manuscript, S.J. supervised the computational work, A.d.S conceived the idea and supervised the project.

