## Supplementary Information for "RNetDys: identification of disease-related impaired regulatory interactions due to SNPs"

**Supplementary Methods**

**Supplementary References**

**Supplementary Figures:**

Fig. S1. Strategy to compute the sign of the regulatory interactions.

Fig. S2. Comparison of the precision in the identification of gene-disease interactions between eQTL and RNetDys.

Fig. S3. Cell (sub)type specific regulatory impairment in PD.

Fig. S4. Cell (sub)type specific regulatory impairment in EPI.

Fig. S5. Cell type specific impairment in T1D.

Fig. S6. Cell type specific impairment in T2D.

Fig. S7. Distribution of the outdegree ratio for specific TFs across cell (sub)types.

Fig. S8. Threshold selection to define accessibility of promoter regions.

**Supplementary Tables:**

Table S1. Single cell datasets used for validation and comparison.

Table S2. Collected datasets to generate healthy cell (sub)type GRNs.

Table S3. Matching of the scRNA-seq and scATAC-seq brain datasets.

Table S4. Literature-based validation of the predicted impaired regulatory interactions.

### RNetDys workflow

#### *Cell (sub)type and state specific GRN inference*

The GRN inference part of RNetDys relies on the combination of multi-OMICS data including single cell datasets (scRNA-seq and scATAC-seq) and prior-knowledge (ChIP-seq and GeneHancer). First, a quality control is performed on the scRNA-seq and scATAC-seq in which any rows (gene or peaks) or columns (cells) having a sum of zero is removed from further analyses.

Then, the following steps are computed to infer the cell (sub)type or state specific regulatory interactions:

(1) TF-Genes interactions: First, using the scRNA-seq data, we pre-selected genes conserved at least in 50% of the cells for candidate interactions. Indeed, we consider genes expressed in the majority of the cells to be representative in the specific cell (sub)type. In addition, from the scATAC-seq peaks matrix, coordinates are extracted to identify accessible promoter regions. Notably, a gene promoter region was identified from the ChIP-seq collected from ChIP-Atlas (Oki et al., 2018), using HOMER (Heinz et al., 2010) annotations by filtering peaks related to gene types annotated as protein coding, and defined as a region between 1500bp upstream and 500bp downstream. A promoter is considered as accessible if its gene has been considered as expressed (conserved at least in 50% of the cells) and at least one ATAC peak is overlapping. The overlap between promoter regions and the peaks coordinates was performed using BEDTools (Quinlan and Hall, 2010) with the parameter -f = 0.48 in reciprocal mode (-r). We identified the overlap parameter f = 0.48 as being the one with the highest probability to capture a real cell (sub)type accessible promoter region. The procedure used to select 0.48 is described in "Identification of accessible gene promoter regions" of the Supplementary Methods. Finally, the resulting overlapping between promoter regions and chromatin accessibility allow us to predict the cell (sub)type or state specific TF-Genes interactions.

(2) Enhancer-Promoters interactions: First, we identified open enhancer regions by intersecting the ChIP-seq data and the scATAC peaks coordinates using BEDTools with the parameter -F 1.0 selecting open enhancer if 100% of the region is accessible. Then, we splitted the scATAC peaks matrix such that one matrix contains accessible promoter regions, obtained previously, and the other one accessible enhancer regions. We then computed the correlation between the two matrices, using the Pearson metric with the propagate R package (Andrej-Nikolai Spiess, 2018) that requires few computational resources to perform correlation of large matrices. Z-scores and corresponding p-values using a one-sided test on a normal distribution is performed for each pairwise correlation generated. Then, a Benjamini-Hochberg multiple test correction was performed on the computed p-values. The network was generated by selecting enhancer regions as sources, and promoter regions as targets, filtering the edges such as p-adjusted value < 0.05 and

keeping promoters for which genes were found in the TF-Genes network. Notably, only positive correlation could be find as being significant as a negative correlation between accessibility peaks translate an absence of interaction between enhancers and promoters. We then retrieved the genes corresponding to the promoter regions using the ChIP-seq data used by RNetDys. Finally, the enhancer-promoter correlation network is intersected with all GeneHancer (Fishilevich et al., 2017) reported connections.

(3) TF-Enhancers interactions: First, enhancers present in the Enhancer-Promoter network are selected. They are then intersected with the ChIP-seq data, using bedtool and -F 1.0, such as if 100% of the TF peak fell inside the enhancer region, then this TF is interacting with the enhancer.

All the interactions of the comprehensive network were then signed based on the scRNA-seq dataset using the Pearson correlation metric between TFs and genes. For TF-Genes interactions, the correlation value defined the sign of the interactions such as positive correlations are most likely activation whereas negative ones are most likely repression. Then, signs for Enhancer-Promoter interactions were determined by computing the sum of correlation values for the TFs binding to the enhancer regulating the specific promoter/gene with the correlation corresponding to the TF-gene relationship (Figure S1) such as:

$$corV_{E_a \to G_b} = \sum_{x} corV_{TF_x \to G_b}$$

With corV: correlation value, TF: transcription factor, E: enhancer, G: gene

Finally, signs for TF-Enhancers were computed by summing, for each TF binding of the enhancer, the TF-genes relationship correlation values for each gene/promoter regulated by the enhancer (Figure S1) such as:

$$corV_{TF_a \to E_b} = \sum_{x} corV_{TF_a \to G_x}$$

With corV: correlation value, TF: transcription factor, E: enhancer, G: gene

*Contextualization towards the disease state to identify candidate impaired interactions*

Based on a GRN from a healthy cell (sub)type or state, the regulatory network was contextualized towards the disease condition of interest based on a list of SNPs. First, promoter regions coordinate for which a TF binding site has been identified is retrieved from the ChIP-seq data. Then, provided SNPs are mapped to these regions and enhancer regions of the GRN using bedtool under the condition that the SNP falls exactly inside one of the regions (parameter -F 1). This step allows the identification of candidate impaired regulatory interactions, including TF-genes and enhancer-promoters, for the specific cell (sub)type. Finally, a TF binding affinity analysis is performed on the SNP impacted regions. The fasta sequences for impacted enhancer and promoter regions were retrieved from genome.ucsc.edu accordingly with the genome assembly, 50bp upstream and downstream were selected from the SNP position and the SNP [ref/alt] alleles were added to the sequence. Then, we used PERFECTOS-APE (E. Vorontsov et al., 2015) to perform the TF motif binding affinity analysis for each SNP on each region found to be involved in regulation. Then, using the cell (sub)type specific GRN, TFs that were binding specifically on the impaired promoter or enhancer were retrieved as well as their dysregulated affinity score. Notably, we used PERFECTOS-APE with the following modified parameters: --pvalue-cutoff 0.05 --fold-change-cutoff 2. Finally, we ranked the TFs to prioritize the regulators that are impaired due to SNPs and hence are most likely to play a role in the dysregulations observed in the disease condition. The rank of each TF regulator was computed as follow:

$$Rank_{TF} = RE \times \frac{NG}{RE} \times \left(\sum |AI|_i^r \times \left(MAF_i^r \times \sum MAF^r\right)\right)$$

With RE: number of regulatory elements regulated by the TF, NG: number of downstream genes across RE, AI: binding affinity impairment log2FC, i: SNPs, r: regulatory element.

**Identification of accessible gene promoter regions**

We intersected ChIP-seq peaks related to gene promoter regions with ATAC peaks from scATAC-seq data to identify accessible cell (sub)type promoter regions using bedtool. In order to define the best threshold to use for the overlapping between the ChIP and ATAC peaks, we collected ChIP-seq from ChIP-ATLAS and compiled four human cell line specific ChIP-seq gold standards (BJ, GM12878, H1 ESC and K-562). We then used all the ChIP-seq collected from ChIP-ATLAS (aspecific) and considered a ChIP peak to be a true positive (TP) if it was found in the cell line specific GS and a false positive (FP) if it was not found in the GS. We computed the percentage of overlaps between ATAC peaks and TPs or FPs ChIP-peaks independently. Then, we computed the delta probability distribution such as: ecdf(TPs overlap) - ecdf (FPs overlap), and selected the highest point = 0.48. Indeed, 0.48 corresponded to the reciprocal threshold for which the probability to capture a TP (cell (sub)type specific ChIP peak) was the highest and was used as default by the RNetDys (Figure S8).

**Generation of the cell (sub)type specific GRNs in healthy condition**

We collected scRNA-seq and scATAC-seq data from human pancreas and brain tissues (Table S2). The scRNA-seq datasets were processed using Seurat v4 (Hao et al., 2021) and, the gene expression and peaks matrices for each cell (sub)type were extracted for each tissue using Signac (Stuart et al., 2020). Annotations were used from their original studies for all tissues.

- Pancreas: we performed the peak calling with MACS2 (-q 0.05 --call-summits) for each cell (sub)type and the peak matrices were extracted for the cell (sub)types having a corresponding scRNA-seq matrix by using the FeatureMatrix function provided by Signac. We then used Seurat to extract all the cell (sub)type scRNA-seq matrices.
- Brain: several datasets were collected to match scRNA-seq and scATAC-seq data in order to extract cell (sub)types and states for different brain regions (Table S3). scATAC-seq fragment files were obtained after request to the authors and the general peaks matrix as well as metadata were retrieved from the public repository of their study (Corces et al., 2020). Each brain region-related scATAC-seq cell (sub)types clusters were annotated using Signac and Seurat with their matched scRNA-seq dataset (Table S3), whereas the cell type annotations were kept from the original study (Corces et al., 2020). We performed the peak calling with MACS2 (-q 0.05 --call-summits) for each cell (sub)type in each brain region. The peak matrices were extracted for the cell (sub)types having a corresponding scRNA-seq matrix by using the FeatureMatrix function provided by Signac. We then used Seurat to extract all the cell (sub)type scRNA-seq matrices. First, we processed the frontal cortex data, imputed the dropouts using MAGIC due to the high rate of zeros (van Dijk et al., 2018) and used the annotations provided by the authors to extract the cell (sub)types (Lake et al., 2018). Of note, excitatory subtypes were merged as excitatory neurons and inhibitory ones as inhibitory neurons to match with the scATAC-seq. Then, we extracted

the cell (sub)types of the substantia nigra for healthy patients while keeping the annotations provided by the authors (Smajić et al., 2022).

Each cell (sub)type GRN was generated using the extracted scRNA-seq and scATAC-seq datasets with the GRN inference part of RNetDys using the default parameters.

**GRN inference benchmarking and comparison to state-of-the-art**

We first assessed the performances of RNetDys to capture cell (sub)type specific TF-Gene interactions and compared to state-of-the-art methods including CLR (Zhang et al., 2016), GENIE3 (Huynh-Thu et al., 2010), SCENIC (Aibar et al., 2017), PIDC (Chan et al., 2017) and ppcor (Kim, 2015). All methods were used with default parameters to infer the TF-Genes networks and applied to 20 single cell RNA-seq datasets collected from six human cell lines (A549, Jurkat, K-562, GM12878, H1 ESC, BJ). Of note, only genes expressed at least in 50% of the cells for each scRNA-seq dataset were provided to the methods to be consistent for the comparison with RNetDys. In addition, predicted (un)directed GRNs were formatted to obtain TF-gene networks by filtering the Source (regulator) such that it contains any human TFs or co-TFs reported in Animal TFDB (accessed on the 08/04/2022)(Hu et al., 2019). Notably, due to large computational resources or a running time higher than two days, five networks could not be generated, including scRNA-seq datasets of one K562, one GM12878 and three H1-ESCs. RNetDys was used with default parameters on the 20 scRNA-seq datasets and scATAC-seq datasets retrieved for each of the six human cell lines (Table S1). We benchmarked the inferred networks against cell line specific GS standard networks compiled from the Cistrome database and computed the precision (PPV) and accuracy (F1-score). Of note, more than one network was generated by RNetDys for each scRNA-seq dataset used for other methods, depending on the number of scATAC-seq datasets. We hence computed the median PPV and F1 score over the networks to have one metric by scRNA-seq, as we had for each state-of-the-art method. We then assessed the performances of RNetDys in capturing cell (sub)type specific enhancer-promoter regulatory interactions. State-of-the-art methods used for the TF-Gene benchmarking did not account for enhancers, as they solely relied on scRNA-seq, and hence we performed a comparison using Cicero (Pliner et al., 2018), a widely used strategy to identify co-accessibility between regulatory regions based on scATAC-seq. We applied RNetDys on twelve combinations of scRNA-seq and scATAC-seq datasets for three human cell lines (Table S1) for which we could compile reliable cell line specific gold standard networks from 3DIV database (GM12878, H1 ESC, BJ/IMR90). We used Cicero on the scATAC-seq datasets using default parameters and annotated the enhancer and promoter regions using the ChIP-seq leveraged by RNetDys. Notably, not significance score was provided on the interactions and hence, accordingly with Cicero guideline we selected interactions with a co-accessibility score greater than zero. Finally, we benchmarked the predicted networks against the human cell line specific GS networks to compute the PPV and F1-scores.

**Compilation of the gold standard networks**

We compiled two types of GS networks, both directed, to assess the performances and validate the specificity in identifying cell (sub)type specific regulatory interactions:

(1) TF-Genes GS networks: for each human cell line, we collected high quality ChIP-seq data specific to the cell line from Cistrome (Mei et al., 2017). The highest quality was defined as peak data passing all the quality control available in Cistrome.
(2) Enhancer-promoter GS networks: for each human cell line, we collected Promoter Capture Hi-C data from 3DIV (Yang et al., 2018) database. We then filtered the GS networks to retain enhancers found in GeneHancer and gene promoter regions defined in the ChIP-seq data retrieved from ChIP-Atlas using BEDTools (Quinlan and Hall, 2010).

**Cell (sub)type specific regulatory mechanisms impaired in diseases**

We performed a general study of cell (sub)type specific impairment in diseases by using prior-knowledge SNPs to validate the relevance of the captured interactions. We first collected single nucleotide variants from ClinVar (Landrum et al., 2018) and extracted SNPs such as the SNV was found at least in 1% of the global population (MAF >= 0.01). Of note, MAF scores were retrieved for each SNV using BioMart R package and the 'hsapiens_snp' dataset. Then, we extracted the SNPs for each disease by selecting the ones that have been reported as being related to the disease in ClinVar and, we performed a systematic extraction using regex with the disease name as pattern. Finally, for each cell (sub)type and each disease, we applied RNetDys using the cell (sub)type GRN and the list of SNPs to capture candidate impaired regulatory interactions, TF binding impairment information and the ranked regulators. Notably, SNPs related to AD were mapped to the brain cortex networks whereas SNPs related to PD were mapped to the midbrain networks.

### Supplementary Figures

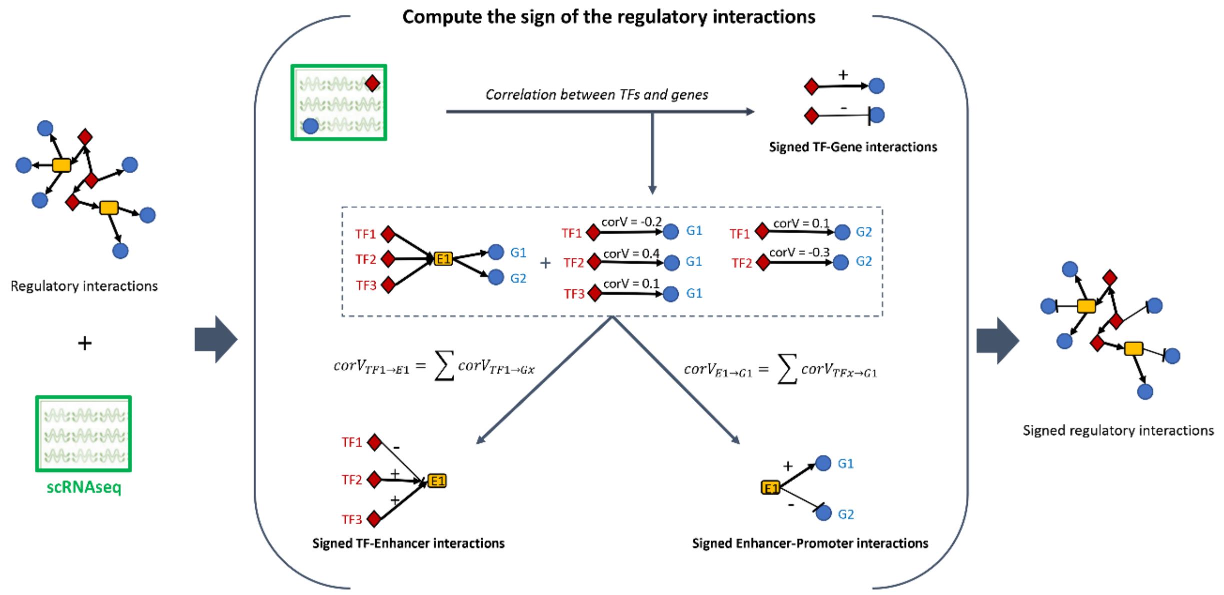


**Fig. S1. Strategy to compute the sign of the regulatory interactions.** The scRNA-seq dataset is used to compute the correlation between the TFs and genes of the GRN. TF-Gene interactions are directly signed using the correlation values. Enhancer-Promoter interactions are signed by summing the correlation values between the TFs binding to the enhancer and the regulated gene/promoter. TF-Enhancer interactions are signed by computing for each TF the sum of the correlation values between the TF and the genes regulated by the enhancer.

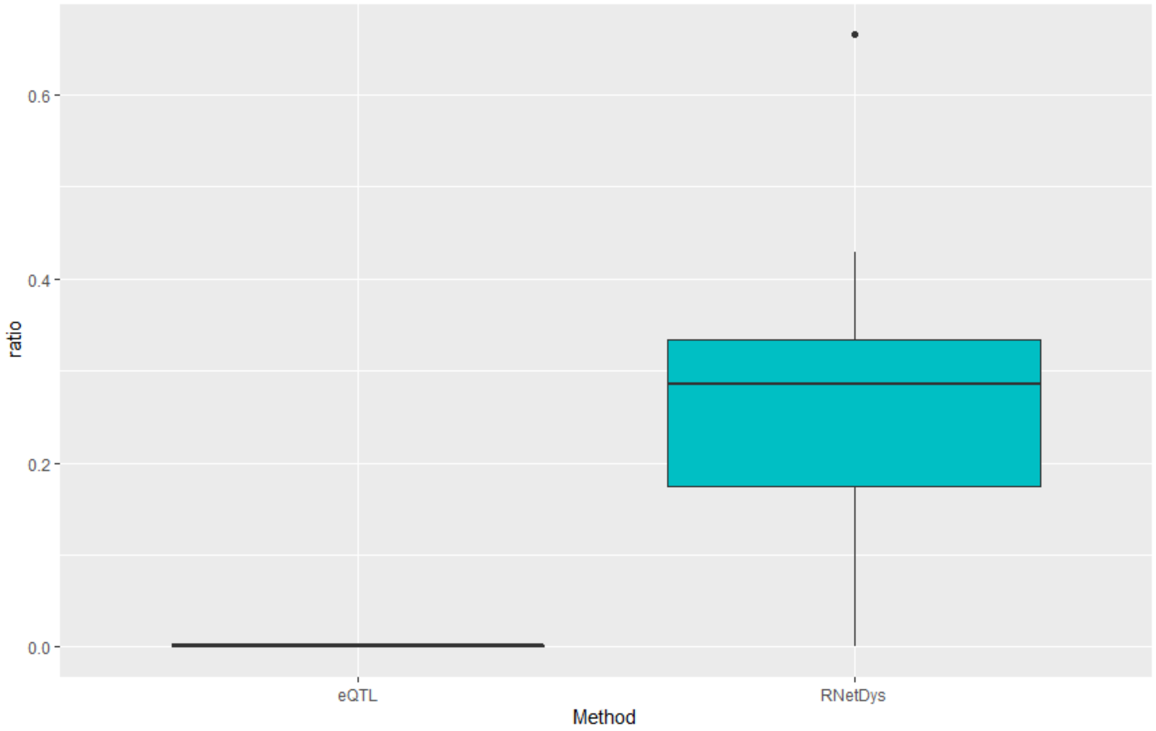


**Fig. S2. Comparison of the precision in the identification of gene-disease interactions between eQTL and RNetDys.** Ratio for the captured genes reported as linked to the disease according to OMIM is represented in y axis. Each boxplot represents ratios across all cell (sub)types for AD, EPI, PD, T1D and T2D.

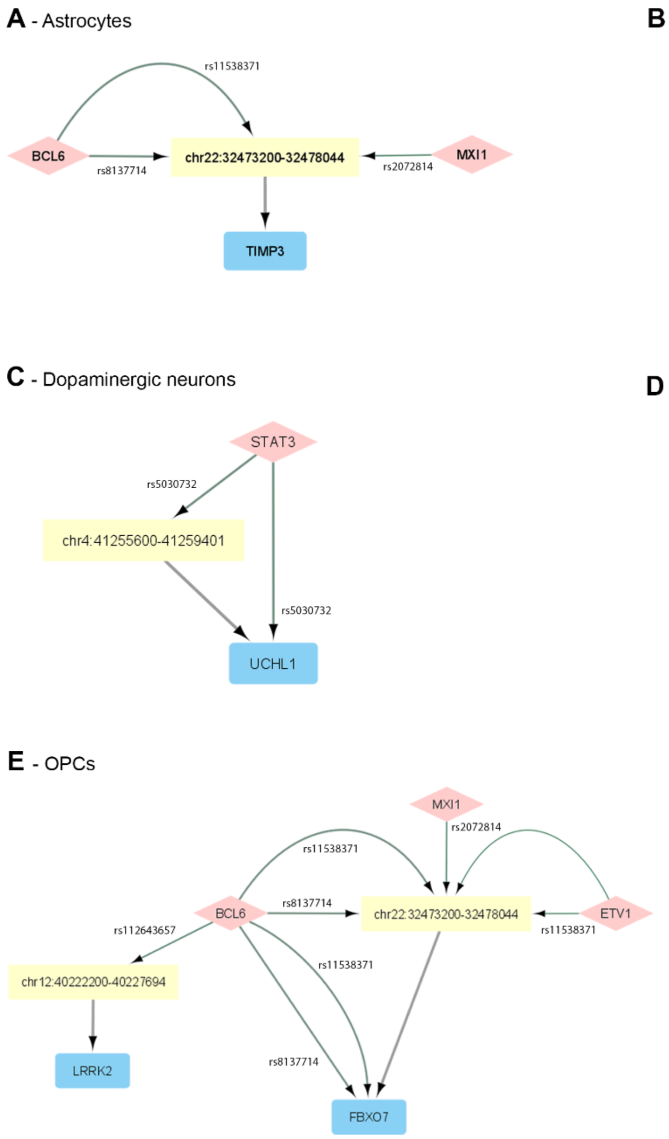


**Fig. S3. Cell (sub)type specific regulatory impairment in PD.** Network visualization of impaired regulatory interactions for (A) astrocytes, (B) excitatory neurons, (C) dopaminergic neurons, (D) oligodendrocytes and (E) OPCs. TFs are represented as diamond in light red, enhancers as yellow rectangles and genes in blue rectangles. Arrows represent activations. The weight of edges from TFs correspond to the strength of the impairment, with the thinnest translating a strong lack of binding affinity and a large edge being a strong increase in binding affinity. The color of the edges from TFs represents the log2FC with green being a decreased affinity and red an increased one.

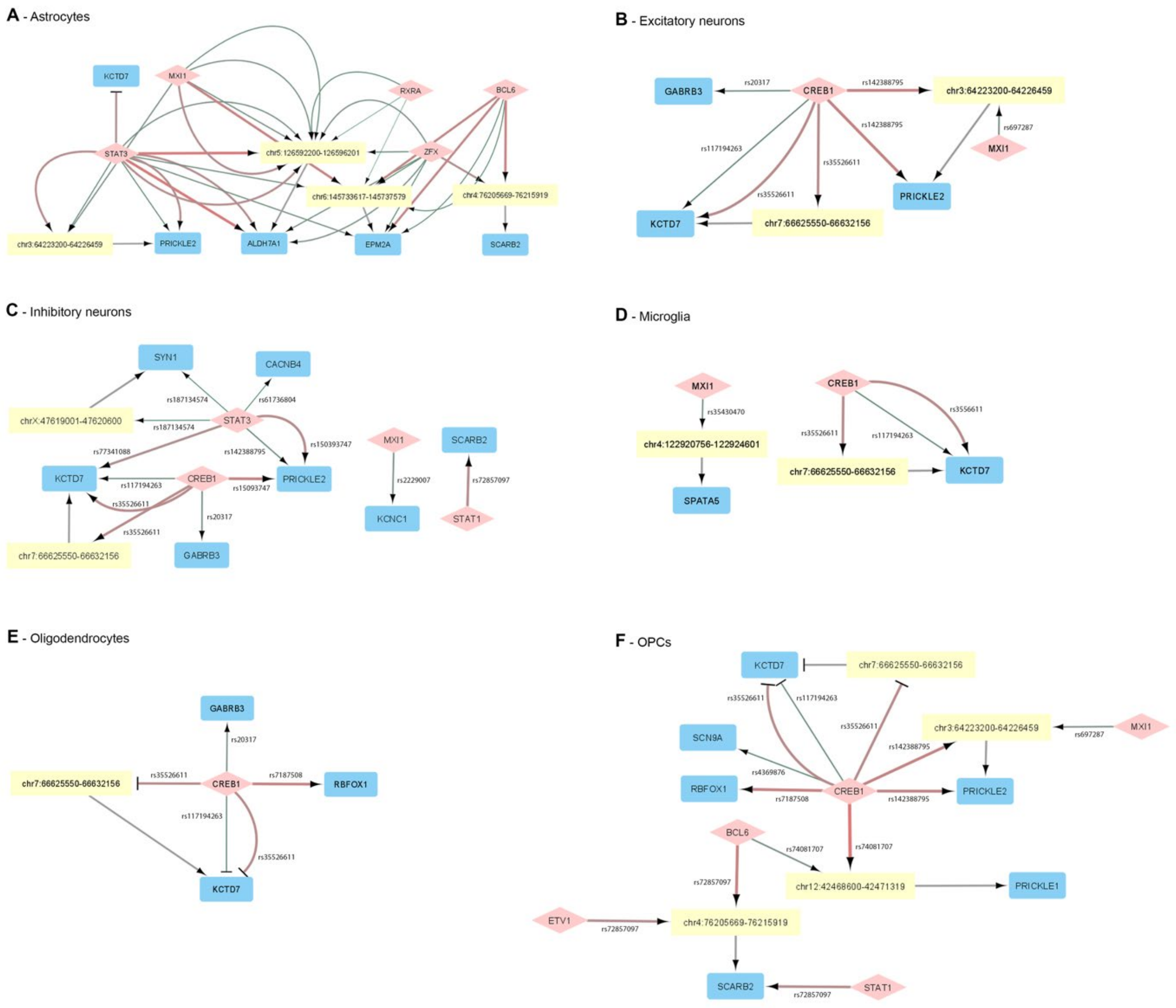


**Fig. S4. Cell (sub)type specific regulatory impairment in EPI.** Network visualization of impaired regulatory interactions for (A) astrocytes, (B) excitatory neurons, (C) inhibitory neurons, (D) microglia, (E) oligodendrocytes and (F) OPCs. TFs are represented as diamond in light red, enhancers as yellow rectangles and genes in blue rectangles. Arrows represent activations and T edges represent repressions. The weight of edges from TFs correspond to the strength of the impairment, with the thinnest translating a strong lack of binding affinity and a large edge being a strong increase in binding affinity. The color of the edges from TFs represents the log2FC with green being a decreased affinity and red an increased one.

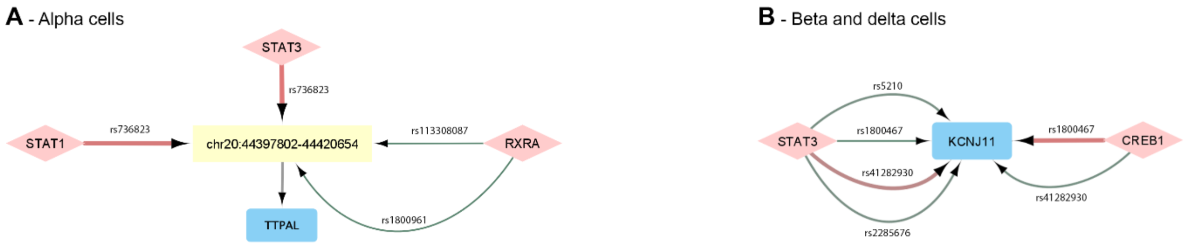


**Fig. S5. Cell type specific impairment in T1D.** Network visualization of impaired regulatory interactions for (A) alpha cells and (B) beta and delta cells. TFs are represented as diamond in light red, enhancers as yellow rectangles and genes in blue rectangles. Arrows represent activations. The weight of edges from TFs correspond to the strength of the impairment, with the thinnest translating a strong lack of binding affinity and a large edge being a strong increase in binding affinity. The color of the edges from TFs represents the log2FC with green being a decreased affinity and red an increased one.

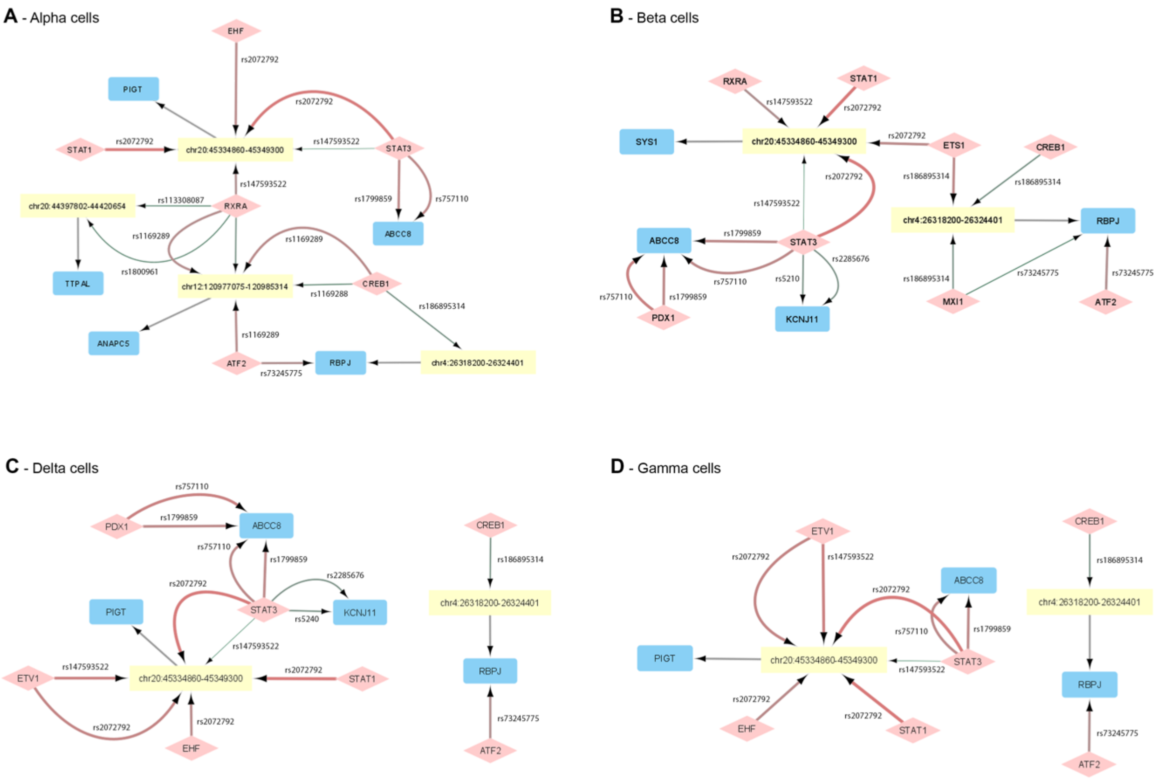


**Fig. S6. Cell type specific impairment in T2D.** Network visualization of impaired regulatory interactions for (A) alpha cells, (B) beta cells, (C) delta cells and (D) gamma cells. TFs are represented as diamond in light red, enhancers as yellow rectangles and genes in blue rectangles. Arrows represent activations. The weight of edges from TFs correspond to the strength of the impairment, with the thinnest translating a strong lack of binding affinity and a large edge being a strong increase in binding affinity. The color of the edges from TFs represents the log2FC with green being a decreased affinity and red an increased one.

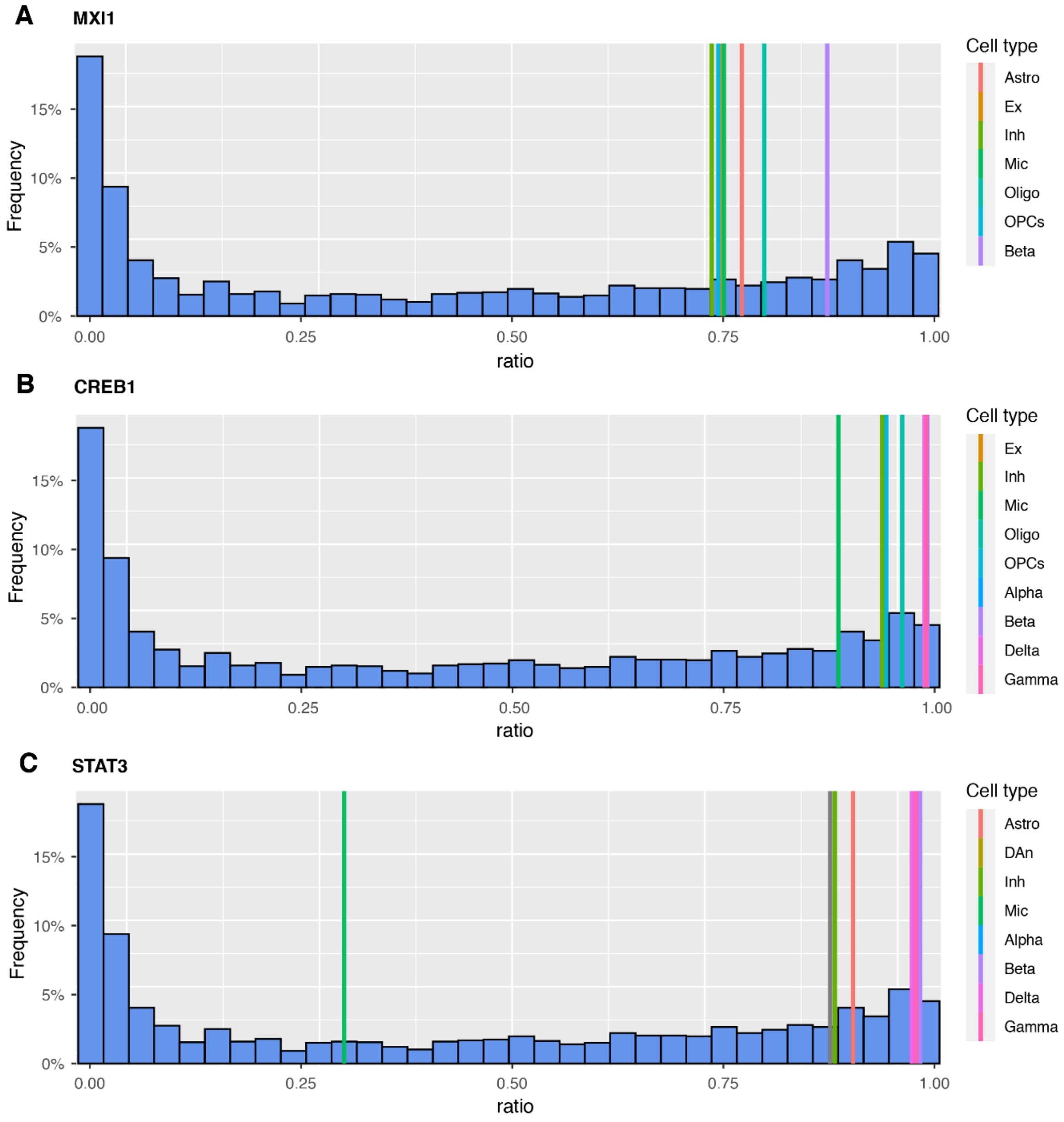


**Fig. S7. Distribution of the outdegree ratio for specific TFs across cell (sub)types.** Histogram showing the frequency of outdegree ratios across all cell (sub)types for three TFs. The outdegree ratio of (A) MXI1, (B) CREB1, and (C) STAT3 in each specific cell (sub)type is represented by coloured vertical lines in the histograms. Astro: astrocytes, Ex: excitatory neurons, DAn: dopaminergic neurons, Inh: inhibitory neurons, Mic: microglia, Oligo: oligodendrocytes, OPCs: oligodendrocyte progenitors, Alpha: alpha cells, Beta: beta cells, Delta: delta cells, Gamma: gamma cells.

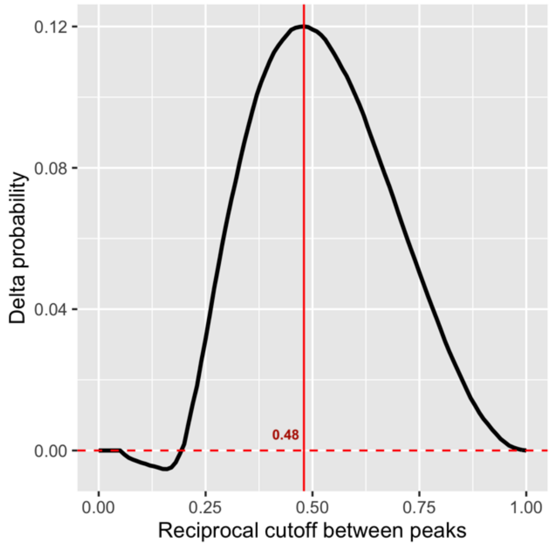


**Fig. S8. Threshold selection to define accessibility of promoter regions.** Delta probability between true positives and false positives. The peak of the distribution, equal to 0.48, corresponds to the highest probability to capture a true accessible promoter region in the cell (sub)type.

### Supplementary Tables

#### Table S1. Single cell datasets used for validation and comparison

| Accession Number | Cell line | Type of data | TF-Promoter benchmarking | Enhancer-Promoter benchmarking |
|---|---|---|---|---|
| GSE100344 | BJ | scRNA-seq | X | X |
| GSE113415 | BJ | scRNA-seq | X | X |
| GSE160910 | BJ | scRNA-seq | X | X |
| GSE166935 | BJ | scRNA-seq | X | X |
| scOpen* | BJ | scATAC-seq | X | X |
| GSE99172 | BJ | scATAC-seq | X | X |
| GSE81861 | GM12878 | scRNA-seq | X | X |
| GSM3596321 | GM12878 | scRNA-seq | X | X |
| GSM4156602 | GM12878 | scRNA-seq | X | X |
| GSM4156603 | GM12878 | scRNA-seq | X | X |
| scOpen* | GM12878 | scATAC-seq | X | X |
| GSE99172 | GM12878 | scATAC-seq | X | X |
| GSE64016 | H1-ESC | scRNA-seq | X | X |
| GSE75748 | H1-ESC | scRNA-seq | X | X |
| GSE81861 | H1-ESC | scRNA-seq | X | X |
| GSM5534158 | H1-ESC | scRNA-seq | X | X |
| scOpen* | H1-ESC | scATAC-seq | X | X |
| GSE99172 | H1-ESC | scATAC-seq | X | X |
| GSE81861 | A549 | scRNA-seq | X | |

| | | | |
|---|---|---|---|
| GSM3271042 | A549 | scRNA-seq | X |
| GSM3271043 | A549 | scATAC-seq | X |
| GSM4224433 | A549 | scATAC-seq | X |
| GSE105451 | Jurkat | scRNA-seq | X |
| 10x platform** | Jurkat | scRNA-seq | X |
| GSE107816 | Jurkat | scATAC-seq | X |
| GSE81861 | K562 | scRNA-seq | X |
| GSE90063 | K562 | scRNA-seq | X |
| GSE113415 | K562 | scRNA-seq | X |
| GSM1599500 | K562 | scRNA-seq | X |
| scOpen* | K562 | scATAC-seq | X |
| GSE99172 | K562 | scATAC-seq | X |

*scOpen: https://github.com/CostaLab/scopen-reproducibility

**10x platform: https://www.10xgenomics.com/resources/datasets/jurkat-cells-1-standard-1-1-0

### Table S2. Collected datasets to generate healthy cell (sub)type GRNs.

| System | Accession | Type of data |
|---|---|---|
| Pancreas | GSE85241 | scRNA-seq |
| | GSM558939 | scATAC-seq |
| Brain | GSE157783 (Healthy) | scRNA-seq |
| | GSE97942 | scRNA-seq |
| | GSE147672 | scATAC-seq |

### Table S3. Matching of the scRNA-seq and scATAC-seq brain datasets.

| scATAC-seq Brain Regions | scRNA-seq Brain Region Matched | Brain region abbreviation |
|---|---|---|
| Substantia Nigra | Human Midbrain (GSE157783, Healthy) | SUNI |
| Middle Frontal Gyrus | Frontal Cortex (GSE97942) | MDFG |

### Table S4. Literature-based validation of the predicted impaired regulatory interactions.

| PD | | | | | | |
|---|---|---|---|---|---|---|
| Source (TF or enhancer) | Gene | RSID | Cell (sub)pop | GWAS | | Cell type specific e-QTL* |
| | | | | SNP Linked to gene | PMID | SNP Linked to gene |
| chr22:32473200-32478044 | TIMP3 | rs11538371 | Astro | | | x |
| chr22:32473200-32478044 | TIMP3 | rs2072814 | Astro | | | x |
| chr22:32473200-32478044 | TIMP3 | rs8137714 | Astro | | | x |

| | | | | | | |
|---|---|---|---|---|---|---|
| chr4:41255600-41259401 | UCHL1 | rs5030732 | DAn | x | 28253266, 25370916, 22839974 | x |
| STAT3 | UCHL1 | rs5030732 | DAn | x | | x |
| NFKB1, STAT3 | PRKAG2 | rs117728810 | DAn | x | | x |
| NFKB1, STAT3 | PRKAG2 | rs66628686 | DAn | x | | x |
| STAT3 | PRKAG2 | rs77902041 | DAn | x | | x |
| chr4:41255600-41259401 | UCHL1 | rs11556273 | Ex | x | | x |
| chr4:41255600-41259401 | UCHL1 | rs5030732 | Ex | x | | x |
| chr4:41255600-41259401 | UCHL1 | rs9321 | Ex | x | | x |
| CREB1 | UCHL1 | rs11556273 | Ex | x | | x |
| chr22:32473200-32478044 | FBXO7 | rs2072814 | Oligo | x | | x |
| chr4:41255600-41259401 | LIMCH1 | rs5030732 | Oligo | | | x |
| chr4:41255600-41259401 | LIMCH1 | rs9321 | Oligo | | | x |
| chr22:32473200-32478044 | FBXO7 | rs11538371 | OPCs | x | | x |
| BCL6 | FBXO7 | rs11538371 | OPCs | x | | x |
| chr22:32473200-32478044 | FBXO7 | rs2072814 | OPCs | x | | x |
| chr22:32473200-32478044 | FBXO7 | rs8137714 | OPCs | x | 18513678 | x |
| BCL6 | FBXO7 | rs8137714 | OPCs | x | | x |
| chr12:40222200-40227694 | LRRK2 | rs112643657 | OPCs | x | | |
| **AD** | | | | | | |
| **Source** | **Target** | **RSID** | **Pop** | **GWAS** | | **Cell type specific e-QTL*** |
| | | | | **Linked to gene** | **PMID** | **Linked to gene** |
| chr14:73135401-73138601 | PSEN1 | rs1800839 | Astro | x | 28821390, 11389157 | x |
| STAT3 | PSEN1 | rs1800839 | Astro | x | | x |
| chr21:26166164-26172001 | APP | rs45476095 | Astro | x | 21654062 | |
| MXI1 | APP | rs45476095 | Astro | x | | |
| chr14:73135401-73138601 | APP | rs459543 | Astro | x | | |
| MXI1 | APP | rs459543 | Astro | x | 21654062, 16685645 | |
| chr14:73135401-73138601 | PSEN1 | rs1800839 | Ex | x | 28821390, 11389157 | x |
| CREB1 | PSEN1 | rs1800839 | Ex | x | | x |
| chr21:26166164-26172001 | APP | rs45476095 | Ex | x | 21654062 | |
| chr21:26166164-26172001 | APP | rs459543 | Ex | x | 21654062, 16685645 | |
| chr14:73135401-73138601 | PSEN1 | rs1800839 | Inh | x | 28821390, 11389157 | |
| CREB1, STAT3 | PSEN1 | rs1800839 | Inh | x | | |
| chr21:26166164-26172001 | APP | rs45476095 | Inh | x | 21654062 | |

| | | | | | | |
|---|---|---|---|---|---|---|
| chr21:26166164-26172001 | APP | rs459543 | Inh | x | 21654062, 16685645 | |
| chr21:26166164-26172001 | APP | rs1800839 | Mic | | | |
| chr21:26166164-26172001 | APP | rs45476095 | Mic | x | 21654062 | |
| chr14:73135401-73138601 | APP | rs459543 | Mic | x | 21654062, 16685645 | |
| CREB1 | PSEN1 | rs1800839 | Oligo | x | 28821390, 11389157 | |
| chr21:26166164-26172001 | APP | rs45476095 | Oligo | x | 21654062 | |
| chr14:73135401-73138601 | APP | rs459543 | Oligo | x | 21654062, 16685645 | |
| chr14:73135401-73138601 | PSEN1 | rs1800839 | OPCs | x | 28821390, 11389157 | x |
| CREB1 | PSEN1 | rs1800839 | OPCs | x | | x |
| chr21:26166164-26172001 | APP | rs45476095 | OPCs | x | | |
| chr21:26166164-26172001 | APP | rs459543 | OPCs | x | | |
| **EPI** | | | | | | |
| **Source** | **Target** | **RSID** | **Pop** | **GWAS** | | **Cell type specific e-QTL*** |
| | | | | **Linked to gene** | **PMID** | **Linked to gene** |
| chr5:126592200-126596201 | ALDH7A1 | rs144272515 | Astro | x | | x |
| ZFX | ALDH7A1 | rs144272515 | Astro | x | | x |
| chr3:64223200-64226459 | PRICKLE2 | rs697287 | Astro | x | | x |
| chr3:64223200-64226459 | PRICKLE2 | rs900641 | Astro | | | |
| chr3:64223200-64226459 | PRICKLE2 | rs142388795 | Astro | x | | |
| STAT3 | PRICKLE2 | rs142388795 | Astro | x | | |
| chr5:126592200-126596201 | ALDH7A1 | rs146562077 | Astro | x | | |
| STAT3 | ALDH7A1 | rs146562077 | Astro | x | | |
| chr3:64223200-64226459 | PRICKLE2 | rs150393747 | Astro | x | | |
| STAT3 | PRICKLE2 | rs150393747 | Astro | x | | |
| chr6:145733617-145737579 | EPM2A | rs2235482 | Astro | x | | |
| BCL6, STAT3, ZFX | EPM2A | rs2235482 | Astro | x | | |
| chr6:145733617-145737579 | EPM2A | rs374338349 | Astro | x | 11735300 | |
| BCL6 | EPM2A | rs374338349 | Astro | x | | |
| chr5:126592200-126596201 | ALDH7A1 | rs60720055 | Astro | x | | |
| chr5:126592200-126596201 | ALDH7A1 | rs72857097 | Astro | | | |
| STAT3 | KCTD7 | rs77341088 | Astro | x | | |

| | | | | | | |
|---|---|---|---|---|---|---|
| chr5:126592200-126596201 | ALDH7A1 | rs900640 | Astro | x | | |
| STAT3 | ALDH7A1 | rs900640 | Astro | x | | |
| ZFX | ALDH7A1 | rs900640 | Astro | x | | |
| chr3:64223200-64226459 | PRICKLE2 | rs697287 | Ex | x | | x |
| CREB1 | GABRB3 | rs20317 | Ex | x | 30074174, 24999380, 25025424 | x |
| CREB1 | KCTD7 | rs117194263 | Ex | x | | |
| chr3:64223200-64226459 | PRICKLE2 | rs142388795 | Ex | x | | |
| CREB1 | PRICKLE2 | rs142388795 | Ex | x | | |
| chr7:66625550-66632156 | KCTD7 | rs35526611 | Ex | x | | |
| CREB1 | KCTD7 | rs35526611 | Ex | x | | |
| CREB1 | GABRB3 | rs20317 | Inh | x | 30074174, 24999380, 25025424 | x |
| CREB1 | KCTD7 | rs117194263 | Inh | x | | |
| CREB1, STAT3 | PRICKLE2 | rs142388795 | Inh | x | | |
| STAT3 | PRICKLE2 | rs150393747 | Inh | x | | |
| MXI1 | KCNC1 | rs2229007 | Inh | x | | |
| chr7:66625550-66632156 | KCTD7 | rs35526611 | Inh | x | | |
| CREB1 | KCTD7 | rs35526611 | Inh | x | | |
| STAT3 | CACNB4 | rs61736804 | Inh | x | | |
| STAT1 | SCARB2 | rs72857097 | Inh | x | | |
| STAT3 | KCTD7 | rs77341088 | Inh | x | | |
| chrX:47619001-47620600 | SYN1 | rs187134574 | Inh | x | | No data on chrX |
| STAT3 | SYN1 | rs187134574 | Inh | x | | No data on chrX |
| CREB1 | KCTD7 | rs117194263 | Mic | x | | |
| chr4:122920756-122924601 | SPATA5 | rs35430470 | Mic | x | | |
| chr7:66625550-66632156 | KCTD7 | rs35526611 | Mic | x | | |
| CREB1 | KCTD7 | rs35526611 | Mic | x | | |
| CREB1 | KCTD7 | rs117194263 | Oligo | x | | |
| CREB1 | GABRB3 | rs20317 | Oligo | x | 30074174, 24999380, 25025424 | |
| chr7:66625550-66632156 | KCTD7 | rs35526611 | Oligo | x | | |
| CREB1 | KCTD7 | rs35526611 | Oligo | x | | |
| CREB1 | RBFOX1 | rs7187508 | Oligo | x | | |
| chr3:64223200-64226459 | PRICKLE2 | rs697287 | OPCs | x | | x |
| CREB1 | KCTD7 | rs117194263 | OPCs | x | | |
| chr3:64223200-64226459 | PRICKLE2 | rs142388795 | OPCs | x | | |

| | | | | | | |
|---|---|---|---|---|---|---|
| CREB1 | PRICKLE2 | rs142388795 | OPCs | x | | |
| chr7:66625550-66632156 | KCTD7 | rs35526611 | OPCs | x | | |
| CREB1 | KCTD7 | rs35526611 | OPCs | x | | |
| CREB1 | SCN9A | rs4369876 | OPCs | x | 23292638, 21698661 | |
| CREB1 | RBFOX1 | rs7187508 | OPCs | x | | |
| chr4:76205669-76215919 | SCARB2 | rs72857097 | OPCs | x | | |
| STAT1 | SCARB2 | rs72857097 | OPCs | x | | |
| chr12:42468600-42471319 | PRICKLE1 | rs74081707 | OPCs | x | | |
| **T1D** | | | | | | |
| **Source** | **Target** | **RSID** | **Pop** | **GWAS** | | **Cell type specific e-QTL** |
| | | | | **Linked to gene** | **PMID** | **Linked to gene** |
| chr20:44397802-44420654 | TTPAL | rs113308087 | Alpha | | | |
| chr20:44397802-44420654 | TTPAL | rs1800961 | Alpha | | | |
| chr20:44397802-44420654 | TTPAL | rs736823 | Alpha | | | |
| CREB1, STAT3 | KCNJ11 | rs1800467 | Beta | x | 25733456, 26937418, 25247988 | |
| STAT3 | KCNJ11 | rs2285676 | Beta | x | 32930968, 29903275, 27249660 | |
| CREB1, STAT3 | KCNJ11 | rs41282930 | Beta | x | 25247988, 22289434, 15115830 | |
| STAT3 | KCNJ11 | rs5210 | Beta | x | 32693412, 33101408, 30641791 | No data |
| CREB1, STAT3 | KCNJ11 | rs1800467 | Delta | x | 25733456, 26937418, 25247988 | |
| STAT3 | KCNJ11 | rs2285676 | Delta | x | 32930968, 29903275, 27249660 | |
| CREB1, STAT3 | KCNJ11 | rs41282930 | Delta | x | 25247988, 22289434, 15115830 | |
| STAT3 | KCNJ11 | rs5210 | Delta | x | 32693412, 33101408, 30641791 | |
| **T2D** | | | | | | |
| **Source** | **Target** | **RSID** | **Pop** | **GWAS** | | **Cell type specific e-QTL** |

| | | | | **Linked to gene** | **PMID** | **Linked to gene** |
|---|---|---|---|---|---|---|
| chr20:44397802-44420654 | TTPAL | rs113308087 | Alpha | | | |
| chr20:44397802-44420654 | TTPAL | rs1169288 | Alpha | | | |
| chr12:120977075-120985314 | ANAPC5 | rs1169289 | Alpha | | | |
| chr20:45334860-45349300 | PIGT | rs147593522 | Alpha | | | |
| STAT3 | ABCC8 | rs1799859 | Alpha | x | 28587604, 26740944 | |
| chr20:44397802-44420654 | TTPAL | rs1800961 | Alpha | | | |
| chr4:26318200-26324401 | RBPJ | rs186895314 | Alpha | x | | |
| chr20:44397802-44420654 | TTPAL | rs2072792 | Alpha | | | No data |
| ATF2 | RBPJ | rs73245775 | Alpha | x | | |
| STAT3 | ABCC8 | rs757110 | Alpha | x | 32660410, 32468916, 32930968 | |
| chr20:45334860-45349300 | SYS1 | rs147593522 | Beta | | | |
| PDX1, STAT3 | ABCC8 | rs1799859 | Beta | x | 28587604, 26740944 | |
| chr4:26318200-26324401 | RBPJ | rs186895314 | Beta | x | | |
| chr20:45334860-45349300 | SYS1 | rs2072792 | Beta | | | |

*https://zenodo.org/record/6104982#.Yq2eUy0RryY
